# The impact of climate and habitat on body shape and size evolution in whip spiders (Amblypygi)

**DOI:** 10.1101/2025.10.06.680801

**Authors:** Paulo Mateus Martins, Gabriela A. Galvão, Diogo B. Provete, Thiago Gonçalves-Souza, Gustavo S. de Miranda

## Abstract

Arthropod body size responds to environmental variation at differing spatial scales. Amblypygi (whip spiders) is an ancient order of arachnids with remarkably conserved body shape, despite their global distribution. While several studies have investigated how body size evolved in spiders, virtually no study has addressed this issue in whip spiders. Here, we analysed how body size and shape of 69 species of Charontid whip spiders evolved in response to climate and habitat. We used Generalized Least Squares to test if bioclimatic variables and habitat influence the variation in body size and shape. Additionally, we fitted alternative macroevolutionary models to body size and shape using Bayesian and Maximum Likelihood approaches. Finally, we estimated phylogenetic signal and tested for differences in evolutionary rates among habitats. Body size decreased slightly with increasing mean annual temperatures and increased with increasing yearly precipitation. Body size evolved following an increasing trend, while the adaptive landscape of body shape seems to have distinct optima, but not rates, for each habitat. Our results support both Cope-Depéret’s and Bergmann’s rule, while habitat had a lesser role. This is the first study to analyze the evolution of Amblypygi phenotypes, which helps in understanding why their morphology is so conserved.

## Introduction

Climate and habitat are important drivers of morphological evolution, particularly when novel environmental characteristics create ecological opportunities that promote adaptive radiation (Mahler et al. 2010; Stroud and Losos 2016; Gillespie et al. 2020). Some of the main influences behind adaptive morphological evolution are a balance between inheritance, selection, and developmental and genetic constraints (Arnold 1992; Freckleton et al. 2002; Blomberg et al. 2003). For example, if divergent or disruptive ecological selection is sufficiently strong, heterogeneous environments may increase morphological diversity by selecting lineages that possess traits (e.g., body size and shape) adapted to diverse niches. This can favour the emergence of specialized adaptations, as subpopulations exhibit high differentiation, which may extend to significant genetic separation (Scheiner et al. 2022). In contrast, selection can favor “jack-of-all-trades” generalist ecomorphs by increasing the fitness of an “average” phenotype that performs equally well in different habitats compared to specialized phenotypes that occupy only a specific portion of the niche space (Remold 2012; Bono et al. 2020; Scheiner et al. 2022).

However, little is known about the processes that lead groups that encounter ecological opportunities at large scales to radiate into phenotypically uniform or distinct species. Importantly, the relative contributions of different climates or habitats in dictating these evolutionary trajectories have been barely investigated with empirical data in different biogeographical regions. In fact, morphological adaptations over evolutionary history are often responses to environmental changes, including historical temperature shifts (Troyer et al. 2022). For example, hotter temperatures are hypothesized to exert selective pressure, favoring smaller body sizes in ectotherms (Horne et al. 2017), because organisms grow faster, but reach maturity at a smaller size. In cold regions, they experience prolonged growth and therefore reach larger body sizes (Angilletta 2004). Additionally, climate can influence body shape, particularly affecting heat exchange and water conservation. For example, sun-exposed spiders have more elongated bodies compared with Sun-protected spiders, as elongated bodies facilitate heat dissipation and reduce surface-area-to-volume ratios (Ferreira-Sousa et al. 2021). However, macroclimate gradients may exert weaker or negligible effects on body size evolution when selection pressures are stronger at finer scales, such as microclimate or microhabitat (Ficetola et al. 2018).

Microhabitats can constrain morphological evolution and favor convergent phenotypic integration even in different regions or climates. Certain habitats, such as shaded or subterranean environments, can buffer organisms from external climatic fluctuations (Villacorta et al. 2008; Scheffers et al. 2014; Keppel et al. 2017). Indeed, previous studies have shown that the effects of climate on morphological evolution are negligible (Mod et al. 2020; Frei et al. 2023), whereas some habitats - particularly caves - can exert strong selective pressures (Howarth 1993; Klaus et al. 2013; Balázs et al. 2021). Caves have limited light input and consequent reduced primary production. For instance, (Balázs et al. 2021) demonstrated that hypogean (cave-dwelling) species are larger than epigean (surface-dwelling) species, even in the same climate. While previous literature suggests that selective pressures from both climate and habitat can influence body size and shape, the relative contributions of these factors to morphological evolution at large macroevolutionary and macroecological scales remain poorly understood. Notably, few studies have investigated the evolution of body size in extant arachnids. However, virtually all comparative analyses have focused on spiders (Ceccarelli et al. 2019; Wolff et al. 2022), and no previous study has analyzed how the morphology of whip spiders (Amblypygi) evolved from a phylogenetic perspective.

The evolution and biogeography of whip spiders offer a unique opportunity to study how macroclimate and microhabitats can shape arachnid morphological diversity over time. Whip spiders split from the most recent common ancestor of Pedipalpi ∼408 myr, during the Lower Devonian, a time in which most continents were around the South Pole (Panotia). Despite this ancient origin, the oldest fossil of Amblypygi closely resembles modern species. Correlations between shield and trunk length/width of modern and fossil species suggest an overlap in the morphological space, highlighting the morphological conservatism of whip spiders over hundreds of millions of years (Haug and Haug 2021). Moreover, genetic studies reveal geographical variation in morphology within the same species, indicating hidden genetic complexity beneath their morphologies (Esposito et al. 2015). For example, there is a widespread sexual dimorphism in whip spiders, and this pattern increases towards the equator, potentially linking intraspecific body size evolution with biogeography (McArthur et al. 2018a). Even though these findings suggest a geographical pattern in body size and shape for some species, as far as we know, there are no studies investigating the ecological factors driving differences in the morphology of whip spiders across biogeographical regions. Indeed, we still do not know whether whip spider species living under tree bark and rocks (epigean) or in caves (hypogean) have similar body size and shape.

Here, we investigate the evolutionary and ecological drivers of body size and shape evolution in 69 species from three genera (54% of all known species of the family) of Charontidae whip spiders occurring on distinct climate regimes and three habitat types. We aim to understand why whip spiders exhibit such remarkable morphological conservatism over hundreds of millions of years, and what processes maintain this stasis despite ecological opportunity. Specifically, we address the following questions: How are habitat and climate related to whip spider morphology? How strong is the phylogenetic signal in whip spider morphological traits? How has whip spider morphology evolved over time? Do different habitats influence the evolutionary rate of morphological traits?

By integrating morphometric, phylogenetic, climatic, and habitat data, we aim to disentangle the relative contributions of climatic and habitat-driven selective pressures and shared evolutionary history in shaping the tempo and mode of morphological evolution in whip spiders.

## Methods

### Organism studied

The order Amblypygi comprises approximately 280 described species (World Amblypygi Catalog 2022; de Miranda et al. 2024). This group has dorso-ventrally flattened bodies, prominent raptorial pedipalps, long antenniform legs, and a lack of silk glands or venom, making them unique among arachnids (Fig. 1). Here, we focused on 69 species from three genera (*Charinus* Simon, 1892, *Sarax* Simon, 1892, and *Weygoldtia* de Miranda, Giupponi, Prendini and Scharff, 2018) of the family Charontidae that were sampled in the most comprehensive phylogeny to date, and to which we have linear traits (de Miranda et al. 2024). This is the most species-rich family within the order (World Amblypygi Catalog 2022). These species inhabit a wide range of tropical and subtropical environments, including caves, forests, and urban areas (see Supplementary Material).

**Figure 1.**
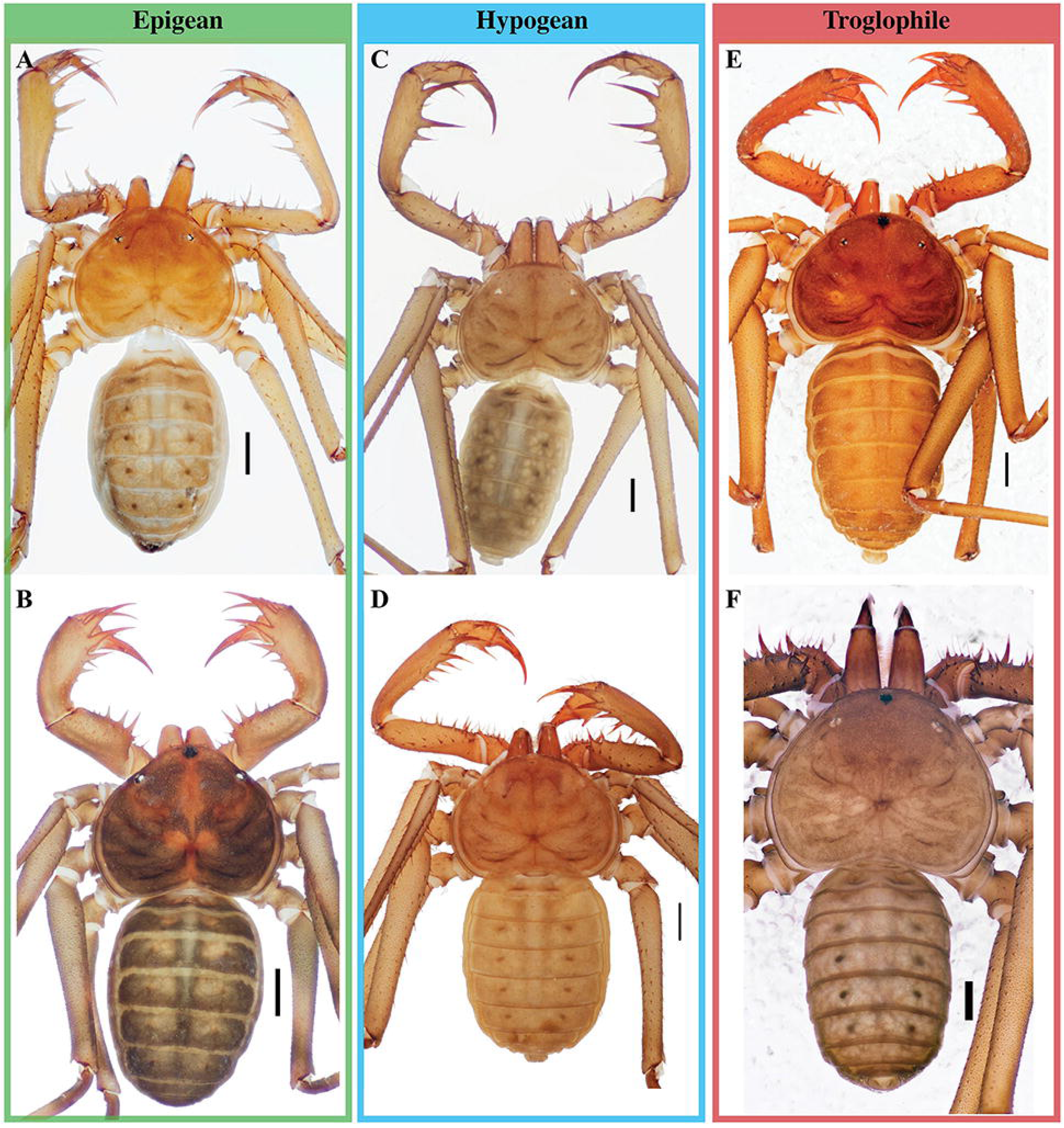
Examples of epigean, hypogean, and troglophile species of whip spiders of the family Charontidae. **A**. Female of *Charinus mocoa* de Miranda et al., 2021, an epigean species from Colombia. **B**. Female of *Sarax gravelyi* de Miranda et al., 2021, an epigean species from Singapore. **C**. Female of *Charinus reddelli* de Miranda et al., 2016a, a hypogean species from Belize. **D**. Female of *Sarax israelensis* de Miranda et al., 2016b, a hypogean species from Israel. **E**. Female of *Charinus brasilianus* Weygoldt, 1972, a troglophile species from the Brazilian Atlantic Forest. **F**. Female of *Charinus apiaca* de Miranda et al., 2021, a troglophile species from the Brazilian Atlantic Forest. Scale bars: 1 mm.

To examine the effects of habitat and climate on body size and shape, we categorized habitat as a factor with three levels, based on data from the literature (Trajano 2012; Trajano and Carvalho 2017): hypogean (n = 24), troglophile (n = 6), and epigean (n = 39). These categories correspond to organisms that primarily inhabit subterranean, surface, and both types of environments, respectively (Figs. 1, 2).

**Figure 2.**
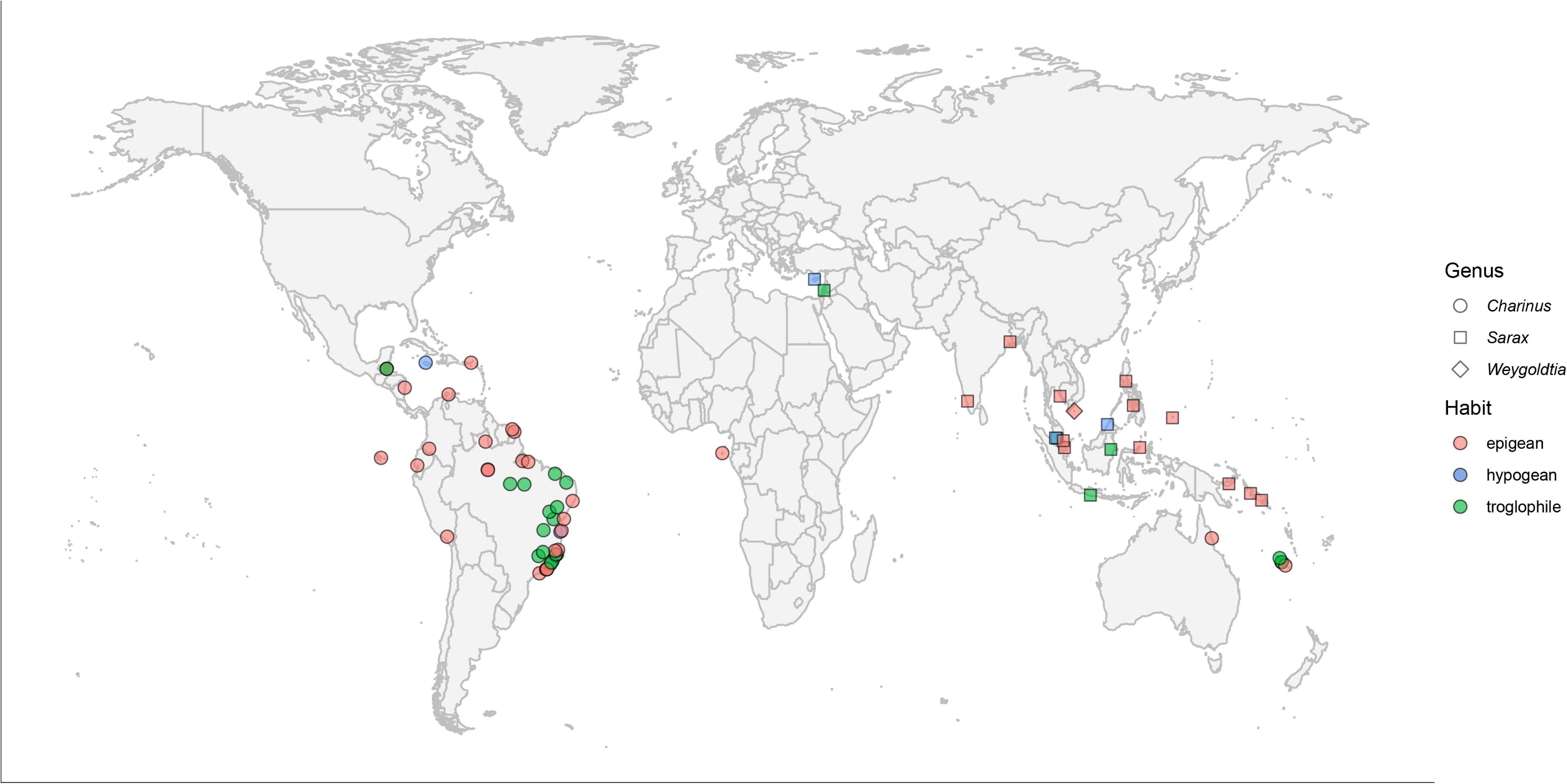
Geographical centroid of the whip spider species used in our study. Colors represent species’ primary habitats. Shapes represent the three Charontidae (former Charinidae) genera for which we had phylogenetic and morphological data.

### Phylogenetic inference and dating

Here, we used a total evidence, dated phylogenetic tree of Amblypygi (Miranda et al. 2021) inferred from a combination of 138 morphological and 3,535 molecular characters (two nuclear: 18S rRNA and 28S rRNA, and three mitochondrial loci: 12S rRNA, 16S rRNA, COI) generated in RAxML-HPC v.8.2.0 (Stamatakis 2014) and dated in BEAST v.1.8.0 (Drummond et al. 2012; De Miranda et al. 2022). Calibration points, models, and further methodological details are described in (De Miranda et al. 2022). This is the largest Amblypygi tree available, including 92 ingroup and 4 outgroup taxa, which represents more than 60% of the Charontidae family and around 35% of the known diversity of Amblypygi (De Miranda et al. 2022; de Miranda et al. 2024).

### Morphometric data

Linear morphometric data were compiled from the literature, including de (Miranda et al. 2021). The average of four linear morphometric measurements (carapace length, carapace width, pedipalp femur length, and pedipalp patella length) was calculated from 193 females from 69 species (see Supplementary Material for the list of species and measurements). Linear measurements of carapace length and width provide a good representation of its shape outline (Réveillion et al. 2022). Only females were used for analysis because whip spiders exhibit strong sexual dimorphism (McArthur et al. 2018a), and several species are known only from female specimens (Miranda et al. 2021). We focused on those four traits because they respond to environmental variation (Weygoldt 2000; McArthur et al. 2018b; Seiter et al. 2022). For example, pedipalp femur length is allometrically related to body size and seems to vary with latitude and climate regimes (McArthur et al. 2018b). Additionally, pedipalp length also plays a role in interference competition, while the total length of walking legs (Réveillion et al. 2022) and pretarsus (Wolff et al. 2015) seem to be related to microhabitat type and rugosity. Concurrently, sexual - instead of natural - selection can also determine the variation in pedipalp length (McLean et al. 2024). Carapace length is a good proxy for body size in whip spiders (Weygoldt 2000; Réveillion et al. 2022).

To obtain a measure of overall body shape, we first calculated the log-shape ratio of linear measurements to obtain size-free shape variables (Mosimann 1970; Claude 2008). Only 64 out of 69 species had data for all four measurements. Thus, body shape analyses have fewer species. First, we computed size as the arithmetic mean of all measurements, then each measurement was divided by size to obtain the shape ratios. Then, we log-transformed this variable to be used in further analyses. Since one degree of freedom is lost during this process, only three variables remain to describe the shape (the last principal component is null).

To summarize the data on body shape, we first performed a phylogenetic Principal Components Analysis (pPCA; (Revell 2009)) on the log-shape ratio variables. The pPCA was performed using a correlation matrix, and traits were assumed to have evolved according to a Brownian Motion (BM) model. The pPCA uses a **C** matrix that describes the variance-covariance between species given the phylogeny and an evolutionary model to calculate the Principal Components (Collyer and Adams 2021). This procedure produced four eigenvectors. To describe body shape, we used all pPCA eigenvectors. This approach avoids the criticism of using only the first eigenvector (Uyeda et al. 2015; Adams and Collyer 2019). Analysis was performed using the R package phytools (Revell 2024).

### Climate data

We examined the relationships between body size, shape, and climatic variables at two scales. The macroclimatic variables included mean annual temperature (BIO1), temperature seasonality (BIO4), annual precipitation (BIO12), and precipitation seasonality (BIO15). First, we used the “micro_global” function from the *NicheMapR* R package (Kearney and Porter 2020) to obtain microclimate data for each locality where specimens were collected. Since whip spiders are predominantly ground-dwelling, inhabiting shaded environments (Weygoldt 2000), we calculated surface temperatures under 90% shade to derive bioclimatic variables using the “calc_biovars” function from the *QBMS* R package (Al-Shamaa et al. 2025). However, microclimate data were not available for six species that occurred on small islands. In these cases, we used WorldClim 2.0 (Fick and Hijmans 2017) bioclimatic data as a substitute. We also ran the models, removing the species for which we imputed WorldClim data to assess how this procedure might affect the results. For both data sets (micro and macroclimate), we calculated the average of each climatic variable for each species, based on the occurrences of collected specimens. The two approaches yielded similar results (Tables 1-2 and S1-S2); therefore, we present only the results for the model with imputed macroclimate data.

**Table 1.** Results of the phylogenetic generalized least squares (pGLS) model examining the relationship between climate and habitat on whip spider body size, accounting for spatial distance and phylogenetic relatedness. The response variable is log-transformed and all continuous predictors are standardized (mean-centered and scaled by standard deviation). Coefficients: Bio 01 = Mean annual precipitation; Bio 04 = Temperature seasonality; Bio 12 = Annual precipitation; Bio 15 = Precipitation seasonality; Habitat: Epigean = occupy primarily above-ground habitats; hypogean = occupy primarily below-ground habitats, including caves; trogrophyle = occupy both above and below ground habitats. “Raw.Estimate” corresponds to the estimates of the analysis of the raw body size as a function of unstandardized estimates.

| Coefficients | Std.Estimate | Standard Error | t | P-value | Raw.Estimate |
| --- | --- | --- | --- | --- | --- |
| Intercept<br>(epigean) | 1.13 | 0.17 | 6.78 | < 0.001 | 3.69 |
| Habitat (hypogean) | 0.18 | 0.060.55 | 3.10 | 0.003 | 0.55 |
| Habitat (troglophile) | 0.01 | 0.08 | 0.16 | 0.88 | -0.04 |
| Bio 01 | -0.09 | 0.03 | -2.64 | 0.01 | -0.05 |
| Bio 04 | 0.04 | 0.05 | 0.91 | 0.37 | 0.002 |
| Bio 12 | 0.10 | 0.04 | 2.5 | 0.02 | 0.00003 |
| Bio 15 | 0.002 | 0.03 | 0.06 | 0.95 | -0.0009 |
| Model fit: Adjusted $R^2 = 0.21$, $F_{(6,57)} = 3.81$, P-value = 0.003 | | | | | |

**Table 2.** Results of the phylogenetic generalized least-squares (pGLS) model testing the effect of climate and habitat on body shape. Columns: F = F-statistic; Z = effect size; PC1.Coef = regression coefficients for the first principal component axis of body shape. Coefficients: Bio 01 = Mean annual precipitation; Bio 04 = Temperature seasonality; Bio 12 = Annual precipitation; Bio 15 = Precipitation seasonality; Habitat: Epigean = above-ground habitats; hypogean = below-ground habitats, including caves; trogrophyle = both above and below ground habitats.

| Coefficients | R<sup>2</sup> | F | Z | P-value | PC1.Coef |
| --- | --- | --- | --- | --- | --- |
| Habitat(Epigean) | 0.05 | 1.6687 | 0.355 | 0.193 | 0.044 |
| Habitat (hypogean) |  |  |  |  | -0.093 |
| Habitat (troglophile) |  |  |  |  | -0.099 |
| Bio 01 | 0.013 | 0.8547 | -0.355 | 0.353 | 0.029 |
| Bio 04 | 0.003 | 0.2055 | -0.701 | 0.738 | -0.051 |
| Bio 12 | 0.057 | 3.7464 | 1.667 | 0.046 | -0.073 |
| Bio 15 | 0.005 | 0.3234 | -0.412 | 0.644 | 0.007 |
| Residuals | 0.870 |  |  |  |  |

## Data analyses

### How are habitat and climate related to whip spider morphology? (question 1)

We used a Generalized Linear Model to test the effect of habitat and climate on body size, which incorporates both phylogenetic and spatial autocorrelation (Freckleton and Jetz 2008). We modelled the log of mean carapace length as a function of habitat, BIO1, BIO4, BIO12, and BIO15. To account for evolutionary relatedness and spatial distance in the residuals, we calculated a phylogenetic variance-covariance matrix based on the most recent phylogeny of Charontidae (De Miranda et al. 2022) and a distance matrix based on species’ mean latitude and longitude. Residual diagnoses were conducted through visual inspection of QQ-plots and residuals vs. fitted plots (Supplementary Material). Residuals showed no deviation from assumptions.

To test for the effect of climate and habit on body shape, we used a multivariate phylogenetic generalized least squares (Adams 2014), as implemented in the R package *geomorph* (Adams and Otárola□Castillo 2013; Baken et al. 2021). Given that this method does not control for spatial autocorrelation, the results did not allow us to evaluate whether the effect of climate on body shape varies geographically. We modelled the phylogenetic principal components (pPCs) of the log-shape ratio as a function of habitat and climate (BIO1, BIO4, BIO12, BIO15). All continuous predictors were mean-centered and scaled by standard deviation. The analysis employed residual randomization and Type I Sum of Squares (SS), following (Adams and Collyer 2018). Residual diagnostics showed no major violations of assumptions.

We ran two models for body size and body shape: one that used WorldClim macroclimate data for the six species for which we could not obtain microclimatic data, and another that included all data points (see Climate data, Tables 1-2, and S1-S2). All continuous predictor variables were standardized by subtracting their values from the mean and dividing by the standard deviation.

### How strong is the phylogenetic signal in whip spider morphological traits? (question 2)

To test whether closely related species have similar body size and shape, we used Blomberg’s *K* statistic (Blomberg et al. 2003; Adams 2014) in the R packages *phytools* (Revell 2024) and *geomorph* (Adams and Otárola□Castillo 2013), respectively.

### How has whip spider morphology evolved over time? (question 3)

To test which model best describes the evolution of whip spider body size, we fit alternative models using Markov chain Monte Carlo (MCMC) in the R package *geiger* (Pennell et al. 2014). To improve model fit (Slater et al. 2012), we used the log of carapace length of the oldest known Amblypygi fossil (*Weygoldtina anglica*, carapace length 3.8 mm, datum from images in (Garwood et al. 2017)), which is at least 313.7 million years old, as a prior at the root. The normal distribution was used as prior, with standard deviation set as 0.1461, which is the log-scaled standard deviation of the species with the most individuals measured in our dataset (*Charinus bonaldoi*). We fitted the following macroevolutionary models: Early Burst (EB), Brownian Motion (BM), Brownian Motion with a direction Trend (Trend), Accelerating-Decelerating with a linear trend (ACDC.lin), Accelerating-Decelerating with an exponential trend (ACDC.exp), and Ornstein-Uhlenbeck with a single stationary peak (SSP). We run three chains with 3,000,000 generations, sampling every 1,000 runs, and discarding the first 30% runs as burn-in. We compared model support using Akaike’s Information Criterion for MCMC samples (AICM; (Raftery et al. 2007)) as implemented by the R package *geiger* (Pennell et al. 2014). Diagnosis of the MCMC chains was performed in the R package *coda* ((Plummer et al. 2006); see Supplementary Material). All models, with the exception of ACDC.lin converged well (Gelman’s PSRF < 1.05, ESS > 200; Supplementary Material).

To determine which model best describes the evolution of body shape, we fitted the Early Burst (EB), Brownian Motion (BM), and single-(OU1) and multi-regime (OUM) Ornstein-Uhlenbeck models to all principal components and compared their fit using the Extended Information Criterion (EIC). In the OU models for body shape, both optima and rates, but not alpha, were allowed to vary among habitats, given the issues with estimating and interpreting the alpha parameter (Cooper et al. 2016). Analyses were conducted in the *mvMORPH* R package (Clavel et al. 2015). Finally, we conducted stochastic character mapping for habitat in the R package *phytools* (Revell 2024) using an all-rates-differ (ARD) model, with 1000 simulations sampling at 10 intervals each.

### Do different habitats influence the evolutionary rate of morphological traits? (question 4)

To test for differences in evolutionary rates (□^2^) of mean body size and shape among habitats, we calculated the net rates of evolution for each habitat (Adams 2014) using the *geomorph* R package (Adams and Otárola□Castillo 2013; Baken et al. 2021). This method calculates the ratio of rates of evolution of a continuous character between groups of species and uses randomization to produce a null distribution (Adams 2014). All data and an R Markdown dynamic document describing the analysis are available in the Supplementary Material.

## Results

The models excluding species with missing data had slightly smaller estimates and greater variability than the model in which we imputed climate data for missing entries. However, both produced comparable results in terms of the direction and magnitude of effects, except for annual precipitation in the body size model (see Tables 1-2 and S1-S2). Below, we present the results for the model with imputed macroclimate data for missing microclimate values.

Our model explained 21% of the variance in whip spider body size (Adjusted R² = 0.21). We found evidence that body size differs among habitats (F = 4.27, df = 2, 57, p = 0.0186), with hypogean species being, on average, 0.55 mm larger than epigean ones (Table 1). Ancestral state estimates for carapace length and habitat (Figure 3) reveal that epigean species had indeed comparatively smaller body sizes. Additionally, we found that mean annual temperature negatively influenced body size, whereas annual precipitation positively influenced body size (Table 1). These translate into body sizes being, on average, around 0.05 mm smaller for each unit of increase in mean annual temperature and 0.003 mm larger for each 100 mm increase in annual precipitation (Table 1). Nonetheless, the precipitation effect on body size disappeared when we compared the models with and without the six species that lacked microclimate precipitation data (see Methods). Therefore, the latter relationship should be taken cautiously.

**Figure 3.**
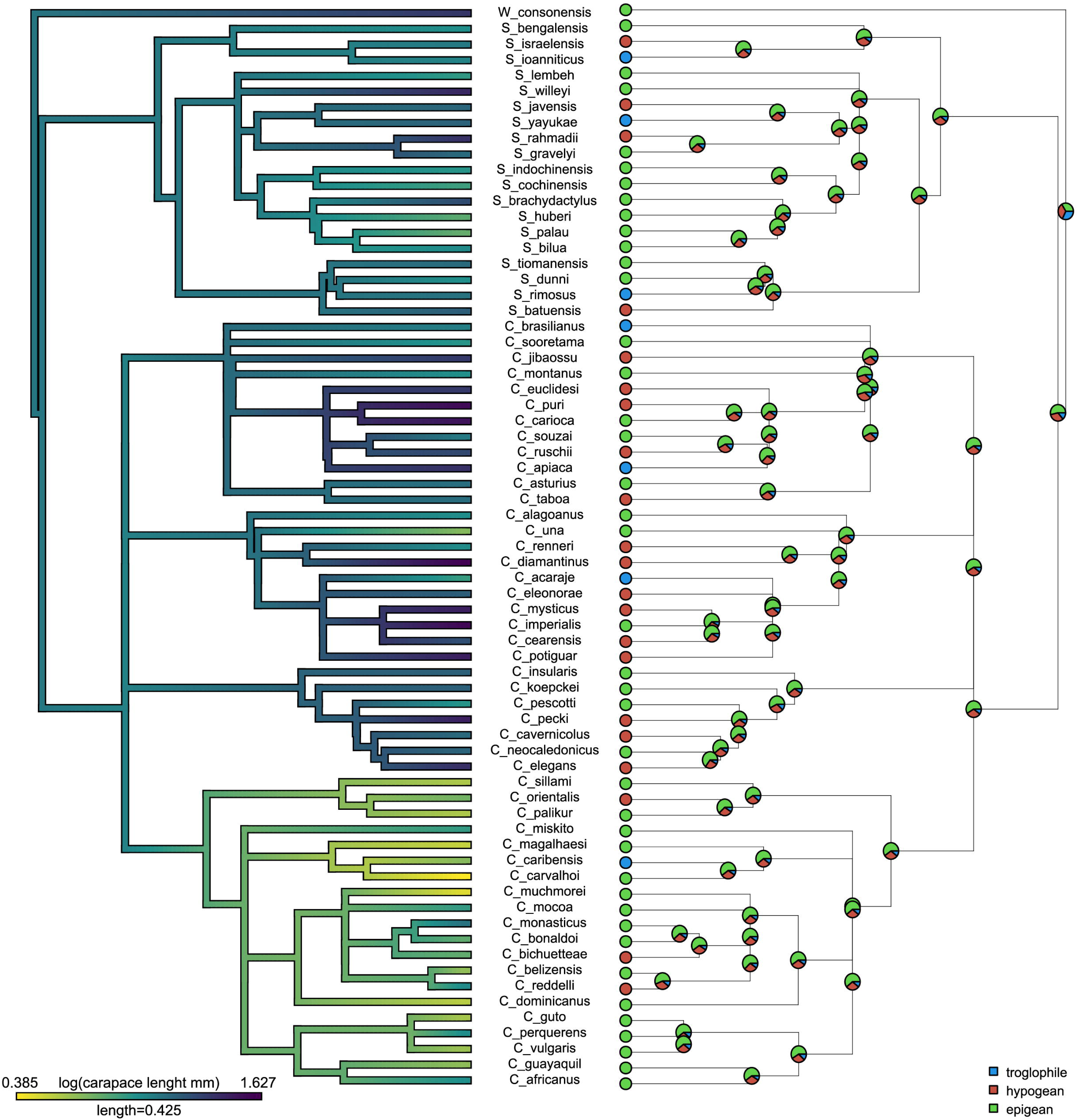
Ancestral state estimations for log-transformed carapace length (left) and habitat (right), estimated using stochastic character mapping. Darker colors indicate larger body sizes.

The first principal component (PC1) of the log-shape ratio accounted for 89.9% of the variance, PC2 for 5.7% (Fig. 4), and PC3 for 4.3%. PC1 contrasts carapace dimensions with pedipalp segment lengths, with high PC1 scores indicating relatively wider/longer carapaces with shorter pedipalps, whereas low PC1 scores indicate proportionally smaller carapaces with elongated pedipalps. PC2 captures shape differences within the carapace and relative pedipalp proportions, with high scores representing wider carapaces paired with longer femurs, and low PC2 scores reflecting more elongated carapaces paired with longer patellae. Annual precipitation was the only variable linked to whip-spider body shape (Table 2), explaining approximately 6% of the variability in the response. This result did not change substantially when we removed from the analysis the six species that lacked microclimate precipitation.

**Figure 4.**
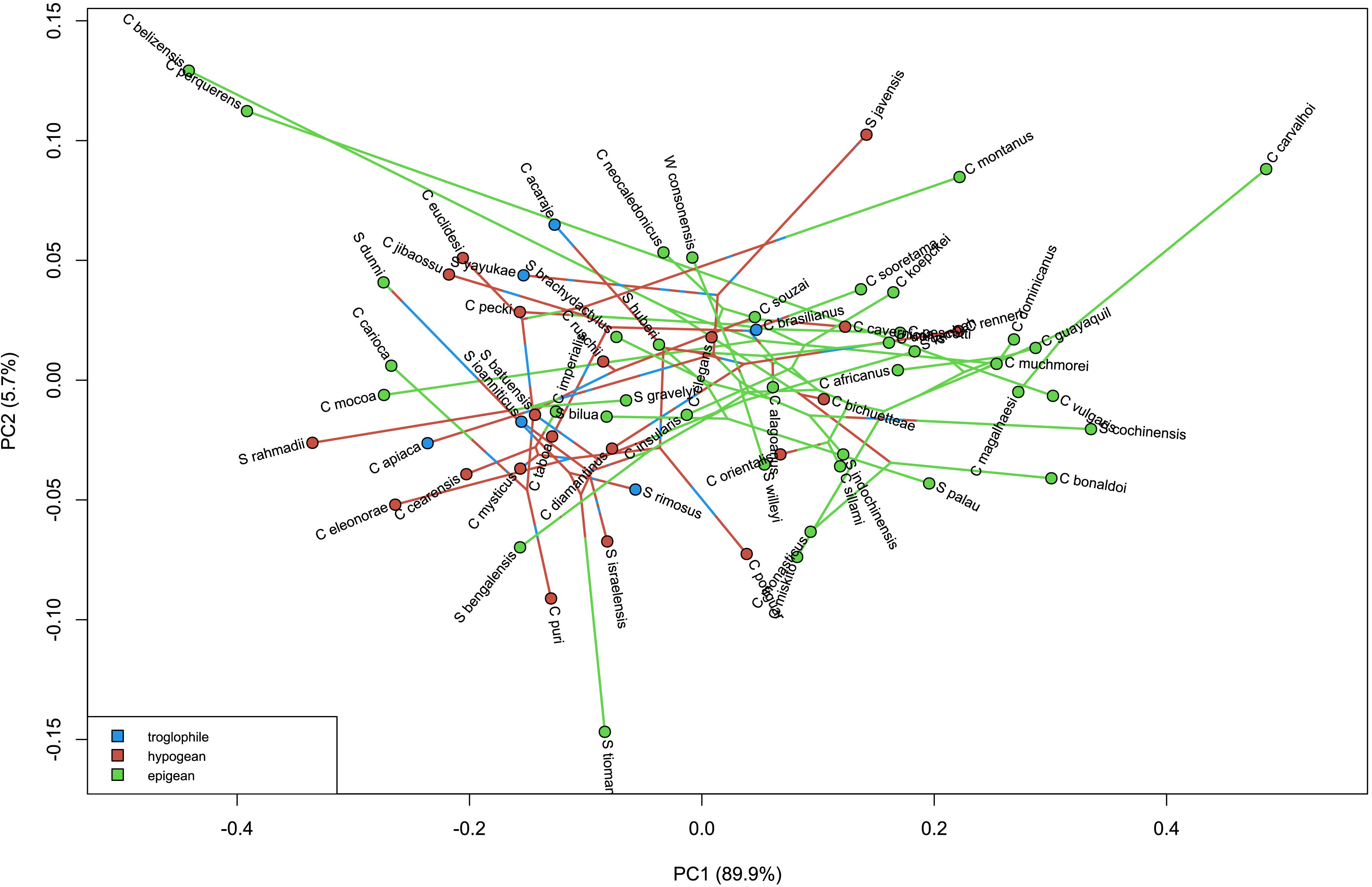
Phylomorphospace plot of the first two phylogenetic principal components of the log-shape ratio, with colors representing different habitats. PC1 contrasts carapace dimensions with pedipalp segment lengths, with high PC1 scores indicating relatively wider/longer carapaces with shorter pedipalps, whereas low PC1 scores indicate proportionally smaller carapaces with elongated pedipalps. PC2 captures shape differences within the carapace and relative pedipalp proportions, with high scores representing wider carapaces paired with longer femurs, and low PC2 scores reflecting more elongated carapaces paired with longer patellae. We estimated the ancestral habitat using the all-rates-differ model (ARD).

We detected a moderate phylogenetic signal for body size (K = 0.6188, *P* = 0.001; Figure 3; Appendix 1), while the phylogenetic signal for body shape was weaker (K = 0.3934, P = 0.185; effect size (Z) = 0.6399, P = 0.257). Both results suggest that body size and shape evolved more slowly, meaning that closely related species resemble each other less than expected under a Brownian Motion model. The small effect size also suggests that the signal is spread across, instead of concentrated in, all dimensions (Adams & Collyer, 2019). For body size, we found strong support for the Brownian motion model with a directional trend (Table 3). In contrast, the multi-peak Ornstein–Uhlenbeck (OUM) model received the strongest support for body shape (Table 4). This result suggests that the optima of body shape, but not size, across species are determined by habitat.

**Table 3.** Comparison of macroevolutionary models for whip spider body size: Early burst (Accelerating-Decelerating) with a linear trend (ACDC.lin), Brownian motion with a directional trend (BM.Trend), Early burst (Accelerating-Decelerating) with an exponential trend (ACDC.exp), Ornstein-Uhlenbeck with a single stationary peak or optimum (OU1), and Brownian motion (BM). dAICM is the difference between the AICM of each model in relation to the best-fitting model. wAICM is a measure of the relative weight of evidence for a given model. Greek letters represent estimated model parameters: evolutionary rate (σ²), linear change in the evolutionary rate (β), directional trend or drift (μ), exponential change in evolutionary rate (r), and attraction towards the optimum trait value (α). Values in parentheses are standard deviations for each parameter.

| Model | dAICM | wAICM | $\sigma^2$ | $\mu$ | $r$ | $\alpha$ | $\beta$ |
| --- | --- | --- | --- | --- | --- | --- | --- |
| BM.Trend | 0 | 0.942 | 0.1661<br>(0.0296) | 0.4207<br>(0.1429) |  |  |  |
| BM | 11.52 | 0.029 | 0.1846<br>(0.0329) |  |  |  |  |
| ACDC.exp | 13.21 | 0.013 | 0.3172<br>(0.2144) |  | -0.4299<br>(0.6925) |  |  |
| OU1 | 13.46 | 0.011 | 0.2435<br>(0.0644) |  |  | 1.281<br>(0.7313) |  |
| ACDC.lin | 14.93 | 0.005 | 0.0931<br>(0.0901) |  |  |  | 7.632<br>(10.44) |

**Table 4.**
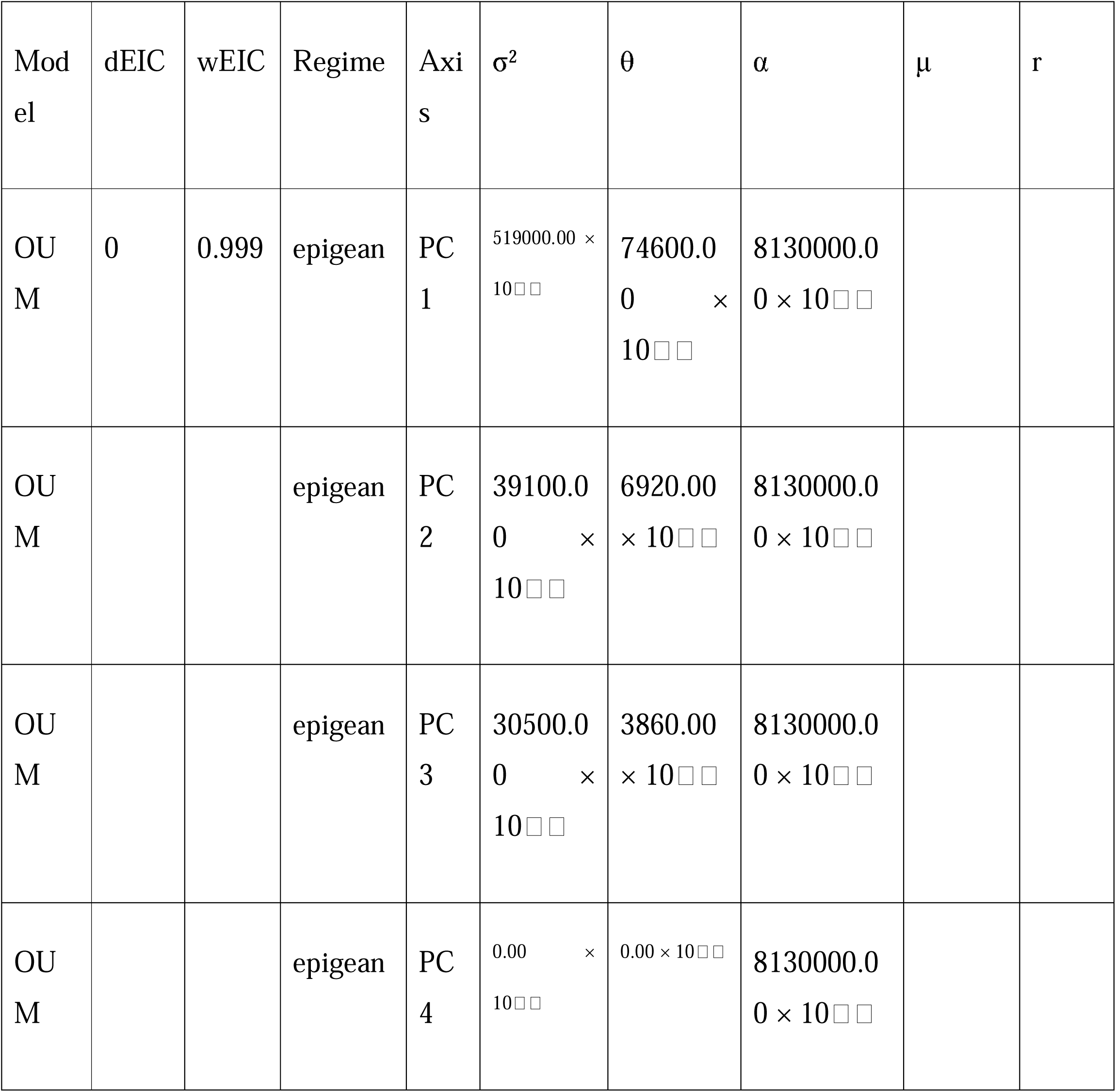
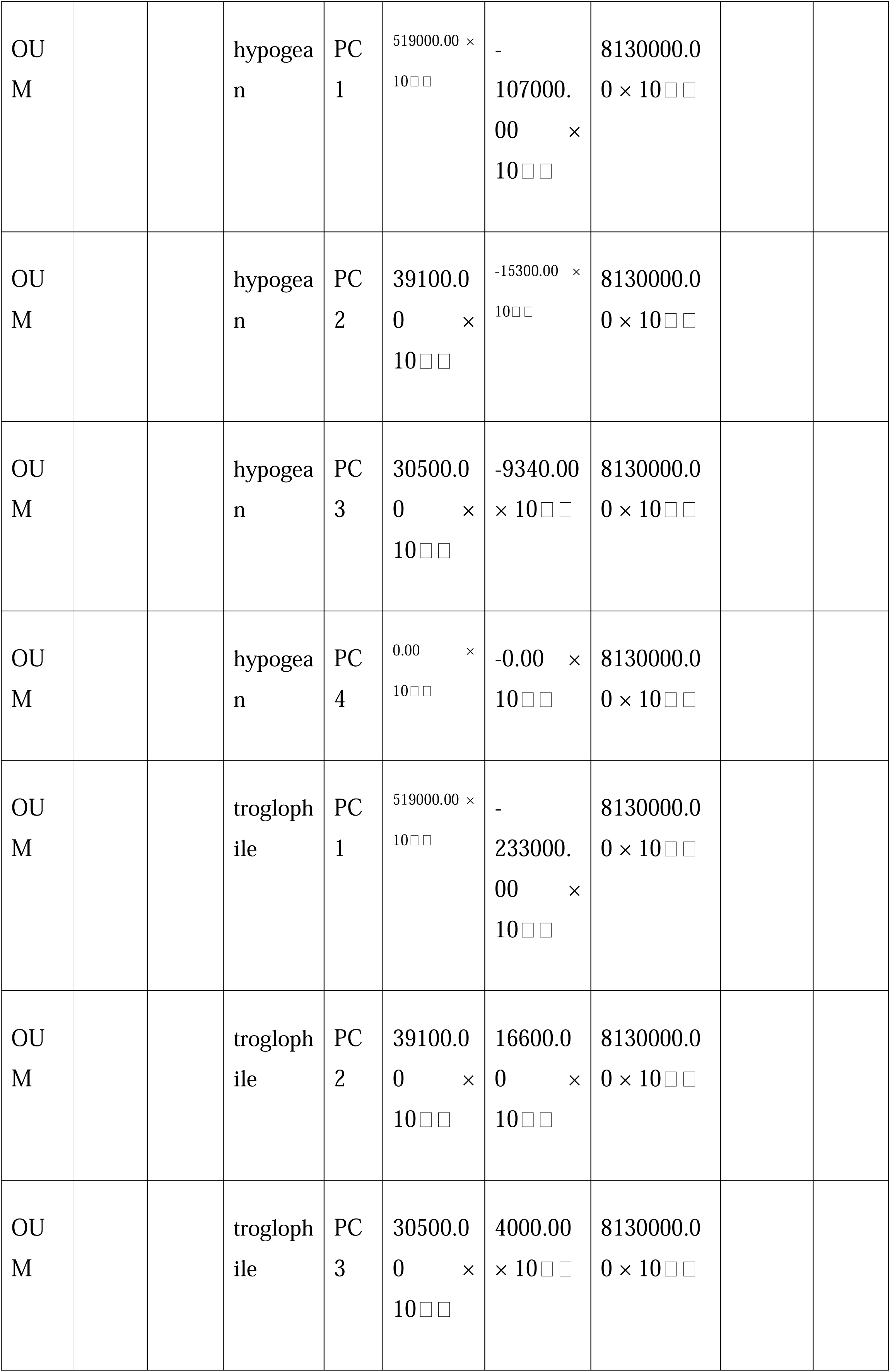
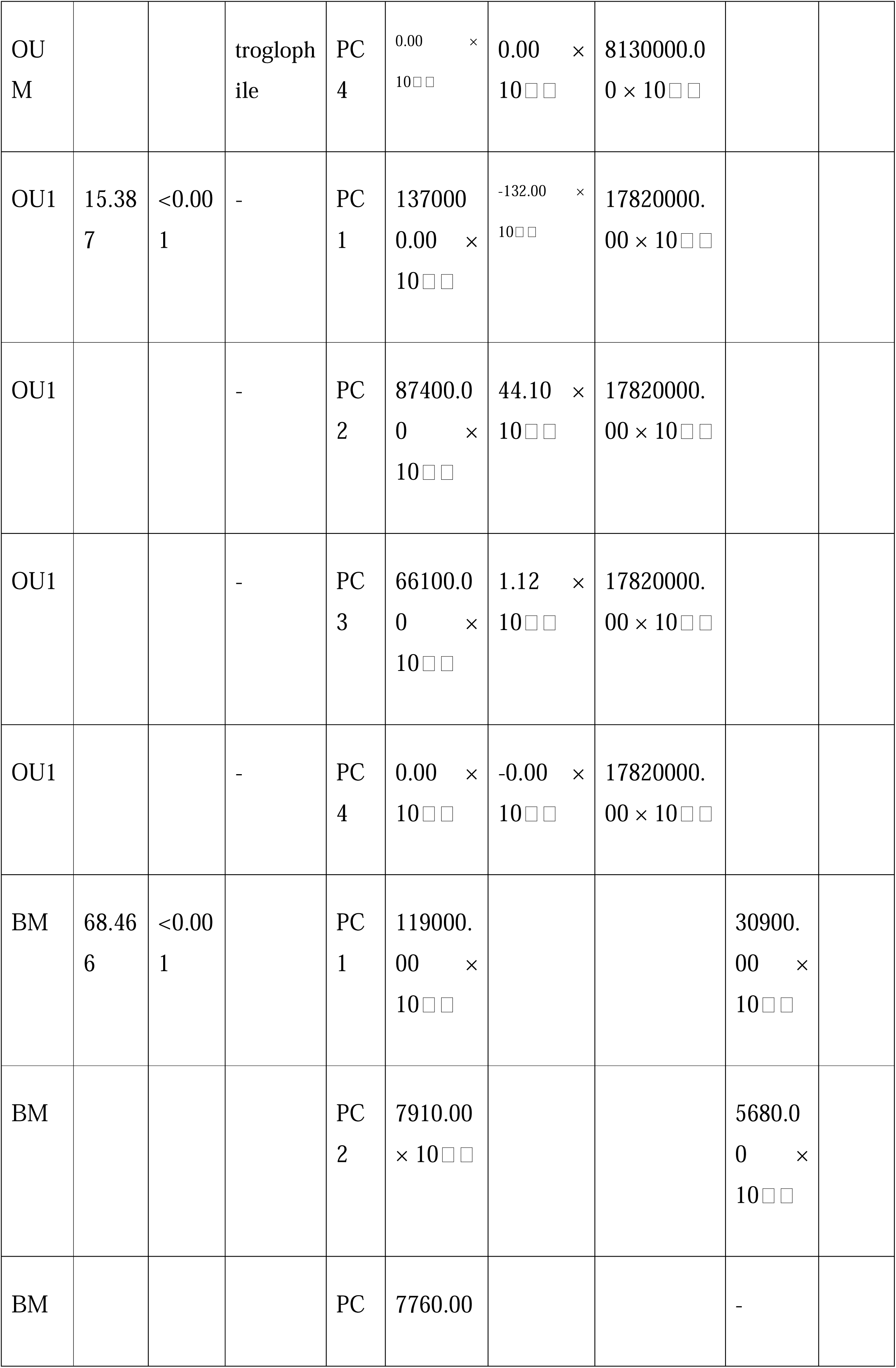
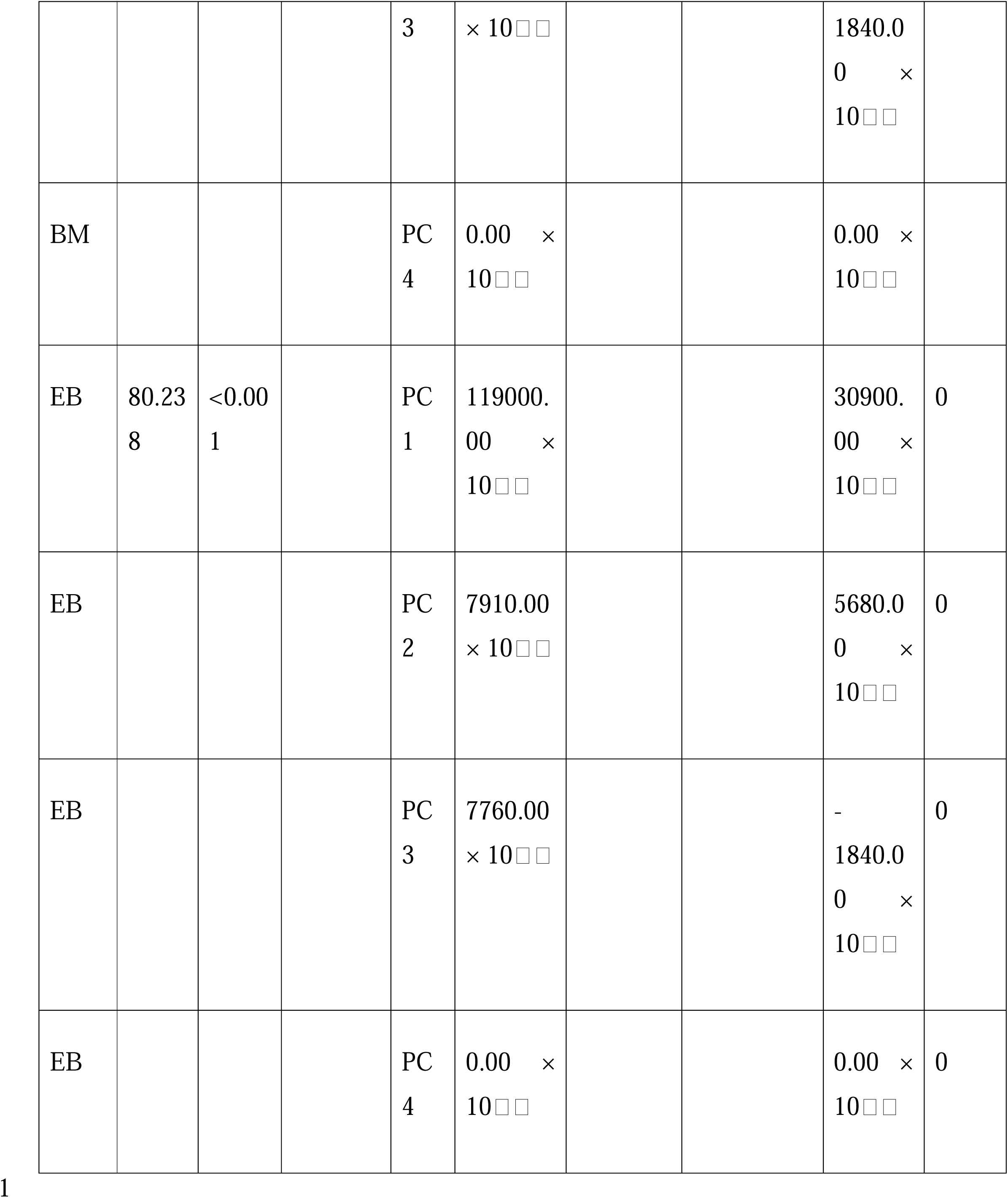
Fit comparison of distinct macroevolutionary models for whip spider body shape: Early burst (Accelerating-Decelerating) (EB), Ornstein-Uhlenbeck with a single stationary peak or optimum (OU1), Ornstein-Uhlenbeck with multiple stationary peaks or optima (OUM), and Brownian motion (BM). dEIC is the difference between the Extended Information Criterion of each model in relation to the best-fitting model. wEIC is the weight of each model based on EIC differences. Greek letters represent estimated model parameters: evolutionary rate (σ²), directional trend or drift (μ), exponential change in evolutionary rate (r), and attraction towards the optimum trait value (α).

Evolutionary rates did not differ among habitats for body size (rate ratio = 2.85, effect size = 0.94, P = 0.19) nor shape (rate ratio = 2.91, effect size = 0.498, P = 0.317). This result mostly agrees with the low phylogenetic signal we found for both traits. However, the best model for body shape was the one in which the optima were allowed to vary by habitat. This suggests that the mode, but not the tempo, of the evolution of body shape is influenced by habitat.

## Discussion

This paper investigated body size and shape evolution based on a unique dataset of a globally distributed group of arthropods, the Amblypygi. Here, we combined data on body size and shape of the largest whip spider family (Charontidae) to test the influence of climate, habitat, and phylogenetic signal in their evolution. We found that body size decreases with increasing mean annual temperatures and increases with increasing yearly precipitation, as well as in hypogean habitats. Additionally, we found a moderate phylogenetic signal in body size, with evidence that this trait followed an increasing trend (positive µ) along the group’s evolutionary history. Together, these findings suggest that body size in whip spiders has been shaped by a combination of environmental influences and gradual evolutionary changes. We detected no phylogenetic signal in body shape, with strong support for a multi-peak Ornstein-Uhlenbeck model, characterized by differing optima and rates per habitat. These findings suggest that selective pressures associated with different habitat types may play a significant role in determining the evolution of whip spider body shape. We also found that annual precipitation had a weak association with body shape. Finally, evolutionary rates did not differ between habitats for either body size or shape. Taken together, our results suggest that multiple adaptive peaks associated with habitats determine whip spider morphology.

The negative association between body size and mean annual temperature agrees with previous studies investigating temperature–size rules in arthropods and supports Bergmann’s rule (Klok and Harrison 2013). The proposed mechanisms for a decreased size in higher temperatures include resistance to starvation and maturation time (Entling et al. 2010). Some authors propose that the association between mean temperature and body size might be an artifact of temperature stability (Cushman et al. 1993). As lower temperatures are negatively correlated with climate stability, larger sizes would permit animals to be more resistant to starvation in colder and unpredictable environments (Cushman et al. 1993). In addition, natural selection seems to favor accelerated maturation of ectotherms in hotter environments, resulting in smaller adult sizes at higher temperatures (Angilletta 2004). Under this hypothesis, accelerated maturation is advantageous for enabling organisms to reproduce sooner, before temperature conditions exceed or approach an organism’s thermal limits (Angilletta 2004).

Furthermore, lower temperatures increase whip spider generation time by approximately two times, compared to those exposed to higher temperatures. This can potentially increase the period of time during which these animals are smaller in size compared to hotter periods (Weygoldt 2000). Temperature and size can also alter the ability of whip spiders to communicate and interact with rivals, potential mates, prey, and predators via their multisensory antenniform legs. Appendages can be modified in larger animals due to thermal inertia, i.e., the amount of time organisms take to heat up and cool down, which tends to be slower in larger animals, and can prevent large individuals from quickly reaching the optimal temperature range for communication (Leith et al. 2021). Given the predominantly tropical origin and distribution of whip spiders and the absence of the effect of temperature seasonality in our models, the second mechanism appears more plausible than the first. Additionally, the primary microhabitats in which whip spiders occur may weaken the direct effect of temperature on body size.

The positive relationship between body size and annual precipitation aligns with the findings of other studies in arthropods (Bidau and Martí 2008; Heino et al. 2019). In the first study, the authors found a positive association between mean annual precipitation and body size of the South American grasshopper, interpreting the relationship as an indirect effect of productivity (Bidau and Martí 2008). In the second study, authors found a positive relationship between body size and precipitation of the driest month, suggesting that water stress during the dry season might also be important for limiting body size (Heino et al. 2019). Precipitation was the only climatic variable associated with body shape in our study. More specifically, whip spiders from wetter environments showed relatively larger carapaces and shorter pedipalps (higher PC1 scores), whereas species from drier environments had relatively longer pedipalps and smaller carapaces. Most whip spider species occupy sheltered microhabitats—such as beneath bark logs and rocks, or within cracks in caves—that may provide similar conditions to those of subterranean environments.

Arthropods have a high surface-to-volume ratio, which presents several challenges for conserving water (Cloudsley-Thompson 1988). The overall expectation is that moisture should have a negative association with body and appendage sizes, as larger sizes lead to smaller surface-to-volume ratios and improved water conservation (Remmert 1981; Entling et al. 2010). A primary strategy arthropods use to avoid water loss is seeking out microhabitats with high humidity (Benoit et al. 2023). Therefore, although our observations contradict the overall expectation of a negative relationship between size and precipitation, the tropical distribution and microhabitats occupied by whip spiders may mitigate the risk of desiccation and selection towards smaller sizes.

Another factor that might help whip spiders retain water is their inherent ability to increase hydrophobicity. Amblypygids have gland openings throughout their body, which secrete a colloidal solution that self-assembles in crystals with high hydrophobic capacity (Wolff et al. 2016). This enables whip spiders to avoid being wet in high-humidity environments and keep moisture inside their bodies. However, the relationship between precipitation and body size is not consistent across animals, as its effect is usually mediated by other factors such as temperature (Sheridan et al. 2022; Fleming and Sheldon 2025) and resource availability (Yom-Tov and Geffen 2006; Gardner et al. 2014).

We found a slight trend toward larger body size in hypogean compared to epigean species. Additionally, habitat had a stronger impact on body shape than on body size. Although not pronounced, the relatively stronger effect of habitat on body shape compared to body size per se likely results from the closer functional connections between shape and ecological factors. Hypogean and troglophile whip spiders have relatively larger carapaces and shorter pedipalps than epigean species. This tendency contradicts the expectation that cave-adapted species evolve troglomorphic traits, such as longer appendages in response to continual darkness and limited resource availability (Culver and Pipan 2019). We propose that the absence of troglomorphic traits in most cave-dwelling species may reflect functional trade-offs unique to the microhabitats occupied by whip spiders in our sample, particularly those from the genera *Charinus* and *Sarax*. Cave-dwelling Charontidae typically inhabit narrow rock crevices during the day and hide in tight spaces when fleeing from potential predators. In these cases, flat bodies may be more advantageous than elongated limbs, potentially affecting the evolution of such traits. Another hypothesis is that most cavernicolous whip spiders did not have enough evolutionary time to develop troglomorphic traits, due to a possible recent colonization of the subsurface. However, the sister species pairs *Sarax ioanniticus* and *S. israelensis* (epigean and hypogean, respectively) have significant differences in middle eye development, despite their recent divergence time (Baken et al. 2021). Comparing multiple cave and non-cave species pairs could elucidate the timing of cave trait changes in whip spiders.

The Brownian-motion model with a positive trend best explained body size evolution. This pattern aligns with Cope-Depéret’s rule, which posits that species tend to increase in size over evolutionary time (Stanley 1973; Heim et al. 2015; Bokma et al. 2016). Advantages associated with larger body size include enhanced prey capture, greater reproductive success, increased metabolic rate (Terblanche et al. 2004) and lower mortality (Stanley 1973; Weygoldt 2000). We also detected a moderate phylogenetic signal for body size and found no differences in evolutionary rates among habitat types. These results suggest that body size evolved more slowly, meaning that closely related species resemble each other less than expected under a Brownian Motion model. This inference is supported by the combination of a moderate phylogenetic signal, similar evolutionary rates across habitats, and a weak indication that cave species are larger than epigean species.

Finally, body shape evolved following a multi-regime Ornstein-Uhlenbeck model, without phylogenetic signal. Together with the absence of rate differences among habitats, these results suggest that the mode, but not the tempo, of the evolution of body shape is influenced by habitat. Thus, stabilizing selection driven by habitat was primordial in body shape evolution, overriding phylogenetic inertia. Although our models underscored a habitat effect on body shape evolution, the phylomorphospace does not appear to show any clear grouping patterns. A previous study on spiders, which analyzed linear measurements separately, also found that OU was the best-fitting model, suggesting that arachnid body shape generally evolves under stabilizing selection (Wolff et al. 2022).

Our results suggest that the body size and body shape of whip spiders evolved under distinct dynamics, with body size exhibiting directional increases through time and only moderate phylogenetic signal. In contrast, body shape evolved under strong stabilizing forces toward different adaptive optima for each habitat. This result reinforces the idea that body shape is more labile than body size. More importantly, our results indicate a jack-of-all-trades strategy in charontid whip spiders, in which a conserved body size confers versatility to occupy distinct habitats. Contrary to predictions of troglomorphic specialization, most cave-inhabiting species did not develop elongate appendages, perhaps indicating that microhabitat-specific functional trade-offs can override generalized cave adaptations for this group of arthropods.

## DATA AVAILABILITY

Scripts used for statistical analyses and figure generation in this study are available in Figshare under the DOI 10.6084/m9.figshare.30287464.

## AUTHOR CONTRIBUTIONS

G.A.G., T.G.S., G.S.M. conceptualized the study. G.A.G., PMM, G.S.M. collated data. P.M.M., G.A.G., D.B.P., T.G.S., G.S.M. critically evaluated the results. P.M.M., G.A.G., D.B.P., G.S.M. ran the molecular phylogenetic and PCM analyses. G.A.G., T.G.S., G.S.M. wrote the first draft. All authors reviewed and edited drafts and approved the final version for publication.

