## Supplementary Material for "The impact of climate and habitat on body shape and size evolution in whip spiders (Amblypygi)"

Scripts used for statistical analyses and figure generation in this study are available in Figshare under the DOI 10.6084/m9.figshare.30287464.

**Table S1.** Results of the phylogenetic generalized least squares (pGLS) model examining the relationship between climate and habitat on whip-spider body size, accounting for spatial distance and phylogenetic relatedness. The analysis excluded species for which we had missing microclimatic data. The response variable is log-transformed and all continuous predictors are standardized (mean-centered and scaled by standard deviation). Bio 01 = Mean annual precipitation; Bio 04 = Temperature seasonality; Bio 12 = Annual precipitation; Bio 15 = Precipitation seasonality; Habitat: Epigean = occupy primarily above-ground habitats; hypogean = occupy primarily below-ground habitats, including caves; trogophyle = occupy both above and below ground habitats. Raw.Estimate = unstandardized estimates.

| **Coefficients** | **Std.Estimate** | **Standard Error** | **T** | **P-value** |
| --- | --- | --- | --- | --- |
| Intercept | 0.97 | 0.13 | 7.29 | < 0.001 |
| Habitat (hypogean) | 0.17 | 0.06 | 2.76 | 0.008 |
| Habitat (troglophile) | 0 | 0.09 | 0.02 | 0.99 |
| Bio 01 | −0.08 | 0.04 | −2.11 | 0.040 |
| Bio 04 | 0.05 | 0.05 | 1.04 | 0.3 |
| Bio 12 | 0.06 | 0.05 | 1.2 | 0.24 |
| Bio 15 | −0.02 | 0.03 | −0.72 | 0.47 |
| Model fit: Adjusted R² = 0.138, F(6, 50) = 2.50, P = 0.034 | | | | |

**Table S2.** Results of the phylogenetic generalized least squares (pGLS) model examining the relationship between climate and habitat on whip-spider body size, accounting for phylogenetic relatedness. This analysis excluded species for which microclimatic data were missing. The response variable represents body shape variation derived from a multivariate shape-space (PCA of log-shape ratios). All continuous predictors were standardized (mean-centered and scaled by standard deviation). Bioclimatic variables: Bio 01 = Mean annual temperature; Bio 04 = Temperature seasonality; Bio 12 = Annual precipitation; Bio 15 = Precipitation seasonality. Habitat categories: Epigean = above-ground species; Hypogean = below-ground (e.g., caves); Troglophile = species occupying both. Model significance was assessed using residual randomization with 1,000 permutations.

| **Predictor** | **Df** | **SS** | **MS** | **R²** | **F** | **Z** | **P-value** |
| --- | --- | --- | --- | --- | --- | --- | --- |
| Bio 01 | 1 | 0.0355 | 0.03552 | 0.00494 | 0.287 | -0.34 | 0.633 |
| Bio 04 | 1 | 0.0685 | 0.06847 | 0.00951 | 0.553 | 0.08 | 0.473 |
| Bio 12 | 1 | 0.3681 | 0.36807 | 0.05115 | 2.975 | 1.39 | 0.092 |
| Bio 15 | 1 | 0.2109 | 0.21087 | 0.0293 | 1.704 | 1 | 0.171 |
| Habitat | 2 | 0.3267 | 0.16334 | 0.0454 | 1.32 | 0.64 | 0.287 |
| **Residuals** | 50 | 6.1863 | 0.12373 | 0.8597 | — | — | — |
| **Total** | 56 | 7.1959 | — | 1 | — | — | — |

|  | **Species** | **Species_phylo** | **N_individuals** | **Mean_Carapace_length** | **SE_Carapace_length** | **Mean_Carapace_width** | **Mean_Pedipald_femur_length** | **Mean_Pedipald_patella_length** | **Habit** | **meanLongitude** | **meanLatitude** | **Realm** | **bio1_wc** | **bio4_wc** | **bio12_wc** | **bio15_wc** | **bio1f** | **bio4f** | **bio12f** | **bio15f** |
| --- | --- | --- | --- | --- | --- | --- | --- | --- | --- | --- | --- | --- | --- | --- | --- | --- | --- | --- | --- | --- |
| 1 | Charinus acaraje | C_acaraje | 4 | 2.54 | 0.19196354 | 3.9875 | 2.6825 | 2.575 | troglophile | -39.391159 | -15.417683 | Neotropical | 23.177208 | 154.665543 | 1373 | 17.0123768 | 21.3464889 | 113.783204 | 1437.33333 | 16.5035431 |
| 2 | Charinus africanus | C_africanus | 6 | 2.76333333 | 0.06669583 | 4.00666667 | 2.02833333 | 2.07166667 | epigean | 6.625353 | 0.285617 | Afrotropical | 21.046648 | 85.4207077 | 2890 | 63.0711632 | 17.1988942 | 92.9936776 | 2070.25 | 54.9687009 |
| 3 | Charinus alagoanus | C_alagoanus | 2 | 2.78 | 0.02 | 3.88 | 2.285 | 2.215 | epigean | -35.898828 | -9.279825 | Neotropical | 23.4847508 | 130.129746 | 1573 | 66.1429977 | 20.6689858 | 87.8534086 | 1416 | 65.5873409 |
| 4 | Charinus apiaca | C_apiaca | 1 | 4.13 | 0 | 5.75 | 4.5 | 4.5 | troglophile | -41.5664 | -21.1538 | Neotropical | 21.9151669 | 223.959641 | 1138 | 66.304863 | 19.1902346 | 207.361515 | 1189 | 63.8209227 |
| 5 | Charinus asturius | C_asturius | 2 | 3.2 | 0.48989795 | 4.55 | NA | 3.45 | epigean | -45.316666 | -23.733334 | Neotropical | 19.1839695 | 249.628388 | 2285 | 43.6113548 | 17.5023493 | 247.516464 | 2078 | 46.6495868 |
| 6 | Charinus belizensis | C_belizensis | 1 | 1.97 | 0 | 2.84 | 3.2 | 2.19 | epigean | -88.68219 | 17.165802 | Panamanian | 23.9085426 | 172.827591 | 1940 | 50.244175 | 20.7086551 | 167.647786 | 1971 | 50.312772 |
| 7 | Charinus bichuetteae | C_bichuetteae | 3 | 2.27333333 | 0.08948929 | 3.2 | 1.77 | 1.77 | hypogean | -53.65806 | -5.878889 | Neotropical | 24.8251762 | 36.8407364 | 1930 | 65.9630508 | 22.3496109 | 48.8220537 | 2140 | 59.9875862 |
| 8 | Charinus bonaldoi | C_bonaldoi | 13 | 2.27923077 | 0.09526254 | 2.99153846 | 1.41307692 | 1.40461539 | epigean | -50.190933 | -1.2795063 | Neotropical | 26.1865215 | 45.6473503 | 2454 | 57.6889153 | 24.0390799 | 34.8759705 | 2312.66667 | 63.8605295 |
| 9 | Charinus brasilianus | C_brasilianus | 5 | 2.978 | 0.05490598 | 4.42 | 2.52 | 2.526 | troglophile | -40.345429 | -19.926747 | Neotropical | 22.5707817 | 199.360855 | 1253 | 51.1029129 | 21.0937647 | 179.430421 | 1250.75 | 50.8461049 |
| 10 | Charinus caribensis | C_caribensis | 4 | 2.0575 | 0.16437254 | NA | NA | 1.5275 | troglophile | -77.65073 | 18.358347 | Panamanian | 24.6036282 | 105.72525 | 1557 | 31.6472607 | 21.5466911 | 120.008379 | 1606.5 | 40.9224908 |
| 11 | Charinus carioca | C_carioca | 1 | 4.69 | 0 | 6.8 | 5.44 | 5.31 | epigean | -43.43533 | -22.933662 | Neotropical | 23.0264893 | 210.863724 | 1373 | 47.4797669 | 19.9080461 | 195.814669 | 1788 | 39.9909081 |
| 12 | Charinus carvalhoi | C_carvalhoi | 1 | 1.47 | 0 | 2.34 | 0.84 | 0.82 | epigean | -60.641003 | 2.587583 | Neotropical | 27.5123329 | 69.2786408 | 1735 | 77.7578812 | 23.4447044 | 41.8904978 | 1677 | 83.9936048 |
| 13 | Charinus cavernicolus | C_cavernicolus | 1 | 3.25 | 0 | 4.7 | 2.56 | 2.47 | hypogean | 165.500055 | -21.500322 | Oceanian | 22.2937088 | 242.662979 | 1495 | 45.1806526 | 19.3743707 | 218.760975 | 1461 | 46.7758305 |
| 14 | Charinus cearensis | C_cearensis | 2 | 3.615 | 0.265 | 5.05 | 3.74 | 3.905 | hypogean | -40.899779 | -3.832944 | Neotropical | 23.6035729 | 75.1637726 | 1276 | 109.5131 | 21.340238 | 82.8575933 | 1124 | 113.550048 |
| 15 | Charinus diamantinus | C_diamantinus | 2 | 5.09 | 0.34063666 | 6.945 | 4.71 | 4.655 | hypogean | -41.322553 | -12.898503 | Neotropical | 22.2495518 | 129.613327 | 941 | 61.5102806 | 19.0785449 | 125.027985 | 1002 | 64.4202552 |
| 16 | Charinus dominicanus | C_dominicanus | 3 | 1.78 | 0.03968627 | 2.52 | 1.20333333 | 1.15333333 | epigean | -71.144615 | 12.015667 | Neotropical | 28.8884831 | 113.645935 | 399 | 91.4447632 | 20.2957022 | 101.52306 | 1426.5 | 60.2743415 |
| 17 | Charinus elegans | C_elegans | 1 | 4.19 | 0 | 5.94 | 3.72 | 3.48 | hypogean | 165.500202 | -21.500265 | Oceanian | 22.2937088 | 242.662979 | 1495 | 45.1806526 | 20.944069 | 210.83959 | 1930 | 47.0902066 |
| 18 | Charinus eleonorae | C_eleonorae | 2 | 3.42 | 0.01632993 | 4.475 | 3.775 | 3.65 | hypogean | -44.169724 | -15.113056 | Neotropical | 22.81773 | 133.218185 | 938 | 100.151245 | 19.1363619 | 190.012813 | 956 | 96.4531083 |
| 19 | Charinus euclidesi | C_euclidesi | 1 | 4 | 0 | 6.06 | 4.55 | 4.25 | hypogean | -42.254444 | -21.937777 | Neotropical | 21.5135937 | 242.60881 | 1255 | 70.9235916 | 18.7458382 | 228.244419 | 1315 | 67.5056899 |
| 20 | Charinus guayaquil | C_guayaquil | 1 | 2 | 0 | 2.75 | 1.33 | 1.23 | epigean | -80.01802 | -2.182331 | Neotropical | 24.9361668 | 106.755653 | 884 | 117.712242 | 21.5610157 | 113.781305 | 793 | 134.195303 |
| 21 | Charinus guto | C_guto | 3 | 1.86666667 | 0.13169831 | 2.47 | NA | NA | epigean | -48.504444 | -1.455833 | Neotropical | 26.4498539 | 41.7467346 | 2652 | 64.9637528 | 24.7561756 | 23.7340516 | 2836 | 57.152866 |
| 22 | Charinus imperialis | C_imperialis | 1 | 4.96 | 0 | 6.88 | 4.9 | 4.75 | epigean | -43.22653 | -22.905758 | Neotropical | 22.8468533 | 208.657181 | 1261 | 49.6736374 | 20.1309199 | 195.355464 | 1629 | 38.2411972 |
| 23 | Charinus insularis | C_insularis | 7 | 3.38 | 0.157918 | 4.59714286 | 2.97857143 | 2.83714286 | epigean | -90.339874 | -0.649724 | Neotropical | 22.7705994 | 166.780243 | 242 | 87.1332169 | 14.7814726 | 142.205849 | 799.333333 | 33.6031099 |
| 24 | Charinus jibaossu | C_jibaossu | 1 | 4 | 0 | 6.1 | 4.56 | 4.38 | hypogean | -45.59651 | -20.28516 | Neotropical | 20.7688026 | 237.519272 | 1391 | 80.4714584 | 17.3875527 | 256.189315 | 1477 | 78.2547298 |
| 25 | Charinus koepckei | C_koepckei | 2 | 3.395 | 0.34468827 | 4.925 | 2.605 | 2.44 | epigean | -71.537361 | -16.409014 | Neotropical | 15.3411875 | 58.4310074 | 59 | 135.873856 | 7.50324786 | 201.451769 | 90 | 141.310097 |
| 26 | Charinus magalhaesi | C_magalhaesi | 1 | 1.72 | 0 | 2.41 | 1.13 | 1.13 | epigean | -59.978424 | -2.975442 | Neotropical | 26.5297394 | 49.0238342 | 2319 | 45.9331131 | 24.3649955 | 28.1633528 | 2233 | 51.1256201 |
| 27 | Charinus miskito | C_miskito | 1 | 2.59 | 0 | 3.52 | 1.92 | 2.16 | epigean | -83.601351 | 13.347422 | Panamanian | 25.5567226 | 82.0507584 | 2317 | 61.7320213 | 25.5567226 | 82.0507584 | 2317 | 61.7320213 |
| 28 | Charinus mocoa | C_mocoa | 1 | 2.6 | 0 | 3.72 | 3 | 2.96 | epigean | -76.651054 | 1.152339 | Neotropical | 23.2355843 | 29.1529121 | 4253 | 23.4834671 | 20.9136905 | 34.3268422 | 4635 | 24.490561 |
| 29 | Charinus monasticus | C_monasticus | 9 | 3.15888889 | 0.16677034 | 4.34666667 | 2.33666667 | 2.61 | epigean | -43.179234 | -22.907164 | Neotropical | 22.8468533 | 208.657181 | 1261 | 49.6736374 | 20.1309088 | 195.35631 | 1629 | 38.2411972 |
| 30 | Charinus montanus | C_montanus | 10 | 2.754 | 0.08315269 | 4.481 | 2.053 | 2.044 | epigean | -40.719585 | -20.131444 | Neotropical | 19.0449791 | 203.835739 | 1326 | 51.4262695 | 18.1080907 | 188.672572 | 1291.75 | 57.6587768 |
| 31 | Charinus muchmorei | C_muchmorei | 3 | 1.59666667 | 0.09322272 | 2.22666667 | 1.08666667 | 1.04333333 | epigean | -64.756355 | 18.352808 | Panamanian | 25.9827728 | 119.273109 | 1185 | 35.1647759 | 25.9827728 | 119.273109 | 1185 | 35.1647759 |
| 32 | Charinus mysticus | C_mysticus | 1 | 4.45 | 0 | 5.94 | 4.45 | 4.31 | hypogean | -42.505817 | -11.43304 | Neotropical | 21.3260841 | 101.247368 | 798 | 88.3270721 | 18.7344065 | 107.241311 | 624.5 | 78.3484439 |
| 33 | Charinus neocaledonicus | C_neocaledonicus | 9 | 3.31555556 | 0.12549974 | 5.20333333 | 3.13 | 3.12444444 | epigean | 166.546144 | -22.171048 | Australian | 21.7470627 | 244.672195 | 1769 | 35.8843155 | 18.5900354 | 220.854605 | 1619.6 | 38.3241921 |
| 34 | Charinus orientalis | C_orientalis | 3 | 2.17333333 | 0.16333333 | 2.94333333 | 1.72666667 | 1.72 | hypogean | -49.633704 | -5.9703693 | Neotropical | 25.5637302 | 53.9879341 | 1793 | 64.5697022 | 23.1033115 | 47.7261553 | 1906 | 75.9602343 |
| 35 | Charinus palikur | C_palikur | 5 | 1.838 | 0.07464081 | 2.426 | NA | NA | epigean | -52.462223 | 4.561417 | Neotropical | 25.5700321 | 41.6121368 | 3194 | 51.3350372 | 24.4571582 | 22.8649585 | 3538.33333 | 50.5423084 |
| 36 | Charinus pecki | C_pecki | 2 | 4.47 | 0.13880442 | 6.48 | 4.75 | 4.41 | hypogean | 164.941104 | -20.688552 | Oceanian | 21.7703972 | 227.782639 | 2030 | 47.9213905 | 18.1619428 | 202.811521 | 1947 | 41.1232381 |
| 37 | Charinus perquerens | C_perquerens | 1 | 2.81 | 0 | 4.81 | 4.06 | 3.85 | epigean | -59.98956 | -3.094908 | Neotropical | 26.8640308 | 52.3582001 | 2323 | 46.3876877 | 24.4939059 | 24.0898947 | 2249 | 52.329691 |
| 38 | Charinus pescotti | C_pescotti | 5 | 2.662 | 0.18073461 | 3.816 | 1.994 | 1.918 | epigean | 145.662484 | -16.719139 | Australian | 22.4147701 | 252.0466 | 1935 | 87.546135 | 20.4725277 | 231.441384 | 1776.75 | 101.242241 |
| 39 | Charinus potiguar | C_potiguar | 1 | 4.5 | 0 | 5.76 | 3.55 | 3.6 | hypogean | -37.688213 | -5.59812 | Neotropical | 27.0392819 | 113.519554 | 776 | 105.602173 | 22.8099627 | 88.4172009 | 775.75 | 104.521321 |
| 40 | Charinus puri | C_puri | 3 | 4.89 | 0.24131584 | 5.96333333 | 4.55 | 4.41666667 | hypogean | -41.940445 | -21.560993 | Neotropical | 22.2426872 | 239.907013 | 1192 | 68.4797211 | 19.5628198 | 229.232925 | 1258 | 66.0137836 |
| 41 | Charinus reddelli | C_reddelli | 3 | 2.84333333 | 0.18223916 | NA | 2.74 | 2.72 | hypogean | -88.730835 | 17.108612 | Panamanian | 23.9085426 | 172.827591 | 1940 | 50.244175 | 20.7086609 | 167.626808 | 1971 | 50.312772 |
| 42 | Charinus renneri | C_renneri | 1 | 2.8 | 0 | 3.95 | 2 | 1.88 | hypogean | -40.32309 | -10.507797 | Neotropical | 23.127594 | 160.687332 | 860 | 35.9685783 | 19.5339293 | 119.311183 | 882 | 34.2479217 |
| 43 | Charinus ruschii | C_ruschii | 9 | 3.75111111 | 0.16426135 | 5.30888889 | 3.62666667 | 3.44666667 | hypogean | -40.601297 | -19.936206 | Neotropical | 19.6889172 | 198.735886 | 1324 | 49.9663315 | 18.4800587 | 182.016106 | 1297 | 56.1821943 |
| 44 | Charinus sillami | C_sillami | 1 | 1.84 | 0 | 2.47 | 1.38 | 1.38 | epigean | -53.047573 | 5.062964 | Neotropical | 25.5176354 | 47.5229797 | 2912 | 46.9966011 | 24.1225917 | 28.0189583 | 2753 | 48.2346813 |
| 45 | Charinus sooretama | C_sooretama | 3 | 2.66666667 | 0.09309493 | 4.04 | 2.08666667 | 2.08333333 | epigean | -40.058975 | -19.036442 | Neotropical | 23.5603542 | 191.119797 | 1195 | 50.149971 | 21.5523479 | 170.690083 | 1202 | 51.3066392 |
| 46 | Charinus souzai | C_souzai | 7 | 3.01 | 0.07758252 | 4.50714286 | 2.56142857 | 2.56714286 | epigean | -40.85554 | -19.22273 | Neotropical | 21.8390732 | 191.417419 | 1208 | 60.4073525 | 19.280784 | 186.478 | 1218 | 66.3903563 |
| 47 | Charinus taboa | C_taboa | 1 | 3.21 | 0 | 4.63 | 3.11 | 3.3 | hypogean | -44.32814 | -19.47491 | Neotropical | 20.9153538 | 201.181595 | 1356 | 87.1231766 | 17.7762475 | 216.313544 | 1435 | 89.9932213 |
| 48 | Charinus una | C_una | 3 | 2.05333333 | 0.02309401 | 2.96 | 1.54333333 | 1.51333333 | epigean | -39.029883 | -15.195411 | Neotropical | 24.1291676 | 146.779694 | 1772 | 10.771246 | 22.1354523 | 105.550594 | 1685 | 9.90192755 |
| 49 | Charinus vulgaris | C_vulgaris | 3 | 1.92666667 | 0.07535103 | 2.59666667 | 1.23666667 | 1.17 | epigean | -38.381575 | -12.875738 | Neotropical | 24.9901218 | 115.631203 | 2003 | 48.3683472 | 23.220277 | 101.830605 | 2111.5 | 54.2013808 |
| 50 | Sarax batuensis | S_batuensis | 3 | 3.41666667 | 0.37834435 | 4.72333333 | 3.43666667 | 3.32 | hypogean | 101.68403 | 3.237831 | Oriental | 25.7548752 | 39.5409508 | 2496 | 27.7766819 | 23.8295554 | 48.9691249 | 2387 | 29.2513728 |
| 51 | Sarax bengalensis | S_bengalensis | 2 | 2.76 | 0.19242809 | 3.525 | 2.68 | 2.64 | epigean | 88.337124 | 22.537171 | Oriental | 26.6495304 | 390.980835 | 1637 | 98.1403351 | 23.2811714 | 386.544054 | 1578.25 | 97.5120244 |
| 52 | Sarax bilua | S_bilua | 2 | 2.825 | 0.125 | 3.775 | 2.685 | 2.475 | epigean | 156.66527 | -7.758767 | Oceanian | 26.1283894 | 42.9318581 | 3588 | 15.2434788 | 24.3938754 | 45.9989264 | 3739 | 15.1414083 |
| 53 | Sarax brachydactylus | S_brachydactylus | 2 | 3.7 | 0.57154761 | 5.325 | 3.565 | 3.39 | epigean | 121.247917 | 14.667068 | Oriental | 24.3687611 | 112.983833 | 2670 | 71.7806778 | 20.5426699 | 107.206597 | 2264.28571 | 59.9534413 |
| 54 | Sarax cochinensis | S_cochinensis | 2 | 2.5 | 0.1 | 3.45 | 1.515 | 1.55 | epigean | 76.3 | 10.683333 | Oriental | 27.3344173 | 147.033569 | 2900 | 103.784493 | 19.6049959 | 119.775529 | 2450 | 107.323274 |
| 55 | Sarax dunni | S_dunni | 1 | 2.75 | 0 | 4.25 | 3.3 | 3.25 | epigean | 159.81181 | -9.131714 | Oceanian | 26.3313732 | 37.7363129 | 2598 | 30.9011421 | 26.3313732 | 37.7363129 | 2598 | 30.9011421 |
| 56 | Sarax gravelyi | S_gravelyi | 2 | 3.225 | 0.025 | 4.38 | 3.025 | 2.815 | epigean | 103.81795 | 1.39075 | Oriental | 26.6542988 | 43.1735878 | 2304 | 22.2945766 | 24.321788 | 66.663281 | 2318 | 18.0942903 |
| 57 | Sarax huberi | S_huberi | 2 | 2.3 | 0.01632993 | 3.4 | 2.11 | 2.12 | epigean | 123.37158 | 9.805584 | Oriental | 25.2851639 | 80.990303 | 2072 | 34.3304329 | 23.2445541 | 70.4556137 | 1685.33333 | 53.3680473 |
| 58 | Sarax indochinensis | S_indochinensis | 2 | 2.68 | 0.06531973 | 3.61 | 2.015 | 2 | epigean | 102.564224 | 11.668076 | Oriental | 26.6860447 | 86.1255264 | 3826 | 92.8027878 | 15.9741609 | 114.804659 | 2367 | 68.8930088 |
| 59 | Sarax ioanniticus | S_ioanniticus | 2 | 2.915 | 0.09389711 | 4.07 | 2.95 | 2.915 | troglophile | 32.796528 | 35.016518 | Saharo-Arabian | 16.8461132 | 646.866028 | 545 | 89.8034821 | 10.2627021 | 469.195096 | 639 | 86.7750543 |
| 60 | Sarax israelensis | S_israelensis | 4 | 3.0375 | 0.17293424 | 4.25 | 2.6725 | 3.05 | hypogean | 35.5504785 | 32.8191725 | Saharo-Arabian | 20.5716457 | 579.312927 | 399 | 108.677368 | 14.9538864 | 470.320736 | 591.666667 | 105.94629 |
| 61 | Sarax javensis | S_javensis | 1 | 3.3 | 0 | 5.25 | 2.75 | 2.5 | hypogean | 111.197773 | -8.123234 | Oriental | 23.4846764 | 49.8595085 | 2341 | 63.9472542 | 23.602325 | 39.8391221 | 4020 | 24.1773535 |
| 62 | Sarax lembeh | S_lembeh | 1 | 2.6 | 0 | 3.68 | 1.91 | 1.84 | epigean | 125.216675 | 1.416694 | Oceanian | 25.4546089 | 25.7559643 | 2781 | 28.0711803 | 23.3545243 | 60.8922829 | 2881 | 37.5043222 |
| 63 | Sarax palau | S_palau | 1 | 2.25 | 0 | 2.9 | 1.56 | 1.5 | epigean | 134.49838 | 7.344517 | Oceanian | 26.1640148 | 55.1956329 | 3526 | 24.3587589 | 26.1640148 | 55.1956329 | 3526 | 24.3587589 |
| 64 | Sarax rahmadi | S_rahmadii | 1 | 4.16 | 0 | 5.92 | 5 | 5.13 | hypogean | 116.98333 | 1.016667 | Oriental | 27.7400417 | 20.2080231 | 2601 | 15.7245131 | 27.7400417 | 20.2080231 | 2601 | 15.7245131 |
| 65 | Sarax rimosus | S_rimosus | 1 | 3.13 | 0 | 4.13 | 2.8 | 2.75 | troglophile | 101.244865 | 3.339183 | Oriental | 27.0089665 | 34.948391 | 2360 | 28.3652687 | 22.9132516 | 117.886178 | 2169.75 | 42.118646 |
| 66 | Sarax tiomanensis | S_tiomanensis | 3 | 3.24333333 | 0.11301426 | 3.67666667 | 2.73333333 | 2.75 | epigean | 103.461078 | 2.783483 | Oriental | 26.6513119 | 61.0537262 | 2929 | 58.8683586 | 26.6513119 | 61.0537262 | 2929 | 58.8683586 |
| 67 | Sarax willeyi | S_willeyi | 1 | 4.31 | 0 | 5.94 | 3.44 | 3.56 | epigean | 150.533986 | -5.844316 | Oceanian | 24.2599487 | 54.1804085 | 4012 | 22.9350414 | 19.3720162 | 56.861212 | 3550.4 | 35.0829607 |
| 68 | Sarax yayukae | S_yayukae | 1 | 3.345 | 0.095 | 4.75 | 3.63 | 3.13 | troglophile | 116.00967 | 6.012445 | Oriental | 27.1182213 | 51.1881561 | 2719 | 34.0643272 | 23.7225837 | 37.2682052 | 3317.5 | 30.8035646 |
| 69 | Weygoldtia consonensis | W_consonensis | 1 | 4.05 | 0 | 6.13 | 3.75 | 3.56 | epigean | 106.595183 | 8.6872 | Oriental | 26.5720062 | 99.8345642 | 2032 | 77.901741 | 26.5720062 | 99.8345642 | 2032 | 77.901741 |
